# Synergistic Screening of Peptide-Based Biotechnological Drug Candidates for Neurodegenerative Diseases using Yeast Display and Phage Display

**DOI:** 10.1101/2023.04.14.536742

**Authors:** Cemile Elif Özçelik, Özge Beğli, Ahmet Hınçer, Recep Erdem Ahan, Mehmet Seçkin Kesici, Talip Serkan Kasırga, Salih Özçubukçu, Urartu Özgür Şafak Şeker

## Abstract

Peptide therapeutics are robust and promising molecules for treating diverse disease conditions. These molecules can be developed from naturally occurring or mimicking native peptides, through rational design and peptide libraries. We developed a new platform for the rapid screening of the peptide therapeutics for disease targets. In the course of the study, we aimed to employ our platform to screen a new generation of peptide therapeutics candidates against aggregation prone protein targets. Two peptide drug candidates for the protein aggregation prone diseases namely Parkinson’s and Alzheimer’s diseases were screened. Currently, there are several therapeutic applications that are only effective in masking or slowing down symptom development. Nonetheless, different approaches are developed for inhibiting amyloid aggregation in the secondary nucleation phase, which is critical for amyloid fibril formation. Instead of targeting secondary nucleated protein structures, we tried to inhibit monomeric amyloid units as a novel approach for halting disease-condition. To achieve this, we combined yeast surface display and phage display library platforms. We expressed α-synuclein, amyloid β_40_, and amyloid β_42_ on yeast surface, and we selected peptides by using phage display library. After iterative biopanning cycles optimized for yeast cells, several peptides were selected for interaction studies. All of the peptides have been used *in vitro* characterization methods which are QCM-D measurement, AFM imaging, and ThT assay, and they have yielded promising results in order to block fibrillization or interact with amyloid units as a sensor molecule candidate. Therefore, peptides are good choice for diverse disease-prone molecule inhibition particularly those inhibiting fibrillization. Additionally, these selected peptides can be used as drugs and sensors to detect disease quickly and halt disease progression.

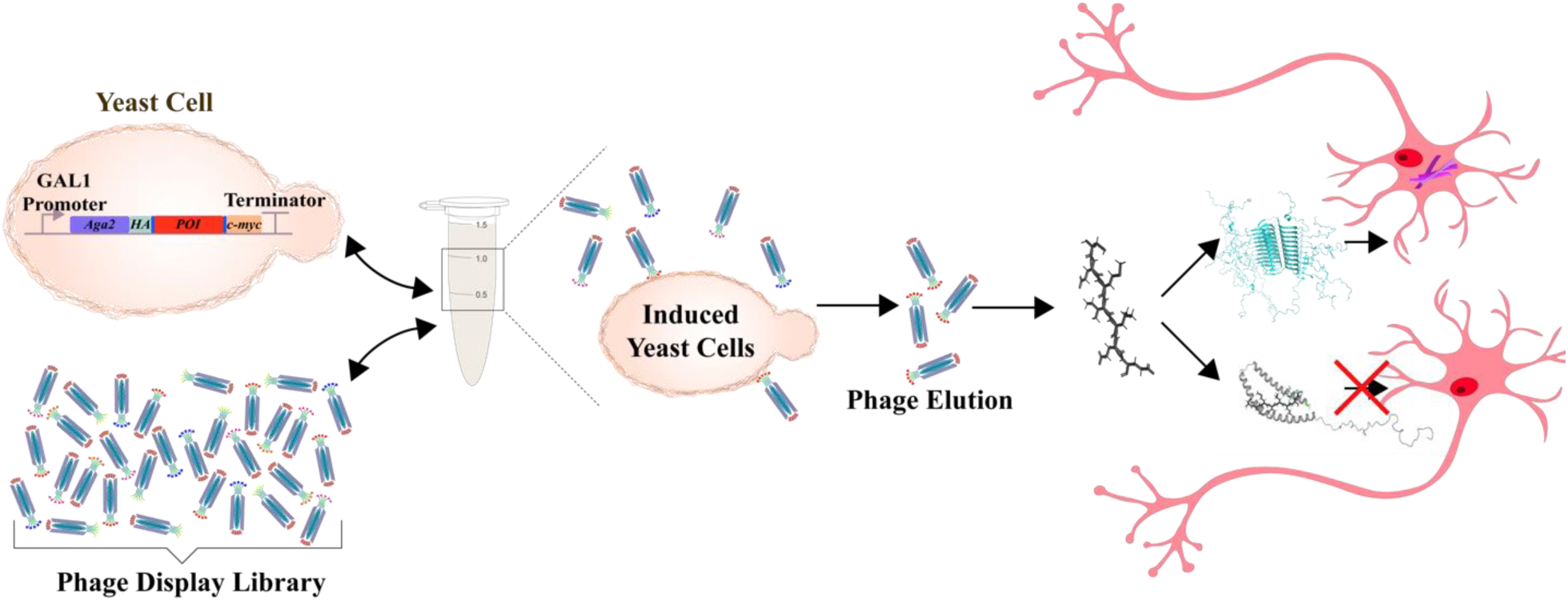

## Introduction

Selection of synthetic peptide-based drug molecules started back in the 80s with the first combinatorial library development, in which foreign DNA pieces were inserted in the phage genome to express random peptide regions in the coat protein of phage ^1^. This approach has provided a tremendous development in the discovery of therapeutic peptides as accumulation of know-how of screening methodologies^2^. Consideration of polypeptides as pharmaceutical agents gained attention with the use of insulin and the synthesis of oxytocin and vasopressin, followed by recombinant insulin production^3–5^. After the potential of therapeutic peptides was revealed, this journey culminated with the approval of more than 80 peptide-based drugs, as well as 400-600 peptides undergoing preclinical trials^6,7^. In light of such peptide-based drug discoveries, display technologies have become the leading technology of peptide selection and screening for pharmaceutical applications.

Using display applications and combinatorial peptide library production, a large and diverse number of peptide pool screenings can be achieved against specific targets, from inorganic compounds to natural products^8–10^. By taking advantage of this and the ease of displaying up to 10^10^ peptide sequences on the coat of the phage, phage display libraries have a wide range of applications, such as the discovery of peptides targeting specific tissues, organs, or tumors, molecular imaging, diagnosis and therapies for neurological disorders, epitope mapping, and novel antibody discoveries, etc.^11–18^. The versatility of peptide screening provides a significant opportunity to select high-affinity bioactive peptides against specific targets^19^. However, for pharmaceutical usage purposes, phage display library-derived peptides need more modifications to delivery, resistance to protease attacks, stability, specificity and clearance as drugs from circulation^20^. Still, phage display is one of the strongest strategies for peptide-drug discovery, which was also appreciated with the one half of the 2018 Nobel Prize in Chemistry “for the phage display of peptides and antibodies” for jumping into a new era for drug molecule discovery by screening combinatorial phage display library^21^.

Alongside the phage display, yeast surface display is an impressive strategy to display a variety of proteins, especially mammalian proteins such as cytokines, surface proteins, antibodies, etc., that undergo post-translational modifications like disulfide bond formation for proper folding and activity, glycosylation as well as presenting the ease of culturing, immobilization and manipulations^22^. Displaying proteins in yeast surface with eukaryotic processing after expression can increase the stability and decrease the vulnerability against temperature, pH, protease attacks, and diverse solvents^23^. By taking advantage of yeast surface display technology as a eukaryotic expression system, many peptides and proteins can be expressed with high stability for protein engineering, antibody, and nanobody screening, as well as fully functionalized enzyme selection^24–28^. Besides, the surface expression of intracellularly toxic proteins is a better approach for interaction studies. Especially in the case of pathogenic amyloids, the monomeric expression outside the yeast cell provides both fully active and proper protein form and decreases the toxic effect compared to the case of residing in the cytosol^29^. Yeast surface display provides immobilization of such dynamic and problematic pathogenic amyloid precursors to discover drug molecules. Till the date, there are various approaches for antibody or drug molecule discovery for such proteins by using yeast surface display as well as phage display separately^30–32^. During the usage of display systems, proteins are immobilized onto a solid surfaces like ELISA plates, polystyrene tubes and magnetic beats, or the screening and panning method requires an additional reporter expression or antibody labelling for sorting the library particles interacted with the target protein^33–38^.

Here, we developed a platform of a combination of phage display and yeast surface display to discover ligand molecules targeting dynamic molecules like amyloid precursors. In our platform, the monomeric versions of α-synuclein, amyloid β_40_, and amyloid β_42_ were expressed on *Saccharomyces cerevisiae* surface as a fusion with the mating factor protein Aga2p to express them in a proper folding with post-translational modifications. Secondly, a phage displayed peptide library is used for the screening of the potential amyloid formation blocking peptides. We have employed precursor proteins related to two different disease conditions. As a proof of concept, we targeted the amyloid formation blocking in Alzheimer’s disease peptides amyloid-β_40_ and amyloid-β_42_ and Parkinson’s disease related α-synuclein protein. Firstly, these proteins, separately displayed on the yeast surface as a fusion of Aga2p. Secondly, for each of the precursor protein cases, M13 phage display library expressing 10^9^ different peptide sequences were introduced to select aggregation-blocking peptides. After the iterative selection of interacting peptides were completed by the biopanning method, peptide sequences were obtained. The peptides were synthetically synthesized and used for further in vitro characterizations such for their amyloid formation blocking potentials. We have carried out amyloid formation inhibition assays using Thioflavin-T staining, monitored the interaction of precursor interaction with pre-formed amyloids with quartz crystal microbalance (QCM). Finally, we observed the amyloid assembly and prevention activities of the inhibitory peptides on the formation of amyloids in real time with atomic force microscopy imaging. In this study, we showed that the combination of yeast and phage display system to form a new generation of peptide-based drug screening platform. The platform provides living yeast cells for protein targets displays, and displayed peptide libraries can be employed sequentially to fast screen of potential peptide-based drug candidates.

## Results and Discussions

### Cell Surface Display of Monomeric Forms of Neurodegenerative Proteins

Regarding neurodegenerative amyloid formation, the monomeric versions of proteins are prone to produce primary nucleation, resulting in ordered amyloid aggregation^39^. We foresee that inhibition of aggregation at this first step is achievable. To do so we aimed to screen and develop peptide inhibitors which can strongly bind to monomeric and oligomeric units of the neurodegenerative proteins. For this reason, we used yeast surface display expressing neurodegenerative proteins (NDPs), which are α-synuclein, amyloid β_40_ and amyloid β_42_, as a fusion with Aga2p (Figure 1A, Supplementary Figure S1A). After NDPs-Aga2p were cloned individually into pETcon vector, and sequences were verified by next-generation sequencing (Supplementary Figure S1B-D) (Addgene plasmid # 41522 ; http://n2t.net/addgene:41522 ; RRID:Addgene_41522). Next, galactose induction was performed to express NDPs in *S.cerevisiae*, and expression characterization was analyzed by immunocytochemistry (ICC). c-Myc tagged NDPs were labelled anti-Myc-tag antibody without any treatment of the cells with membrane solubilizing agents. Fluorescence of the surface displayed NDPs were achieved by detection of Myc-tag antibody with its specific antibody conjugated with DyLight550 in order to indicate the surface display verification and displaying efficiency. Immunostained *S.cerevisiae* cells were examined by fluorescence microscopy to prove the surface display of the target proteins We observed that immunostained cells expressing amyloid β_40_-Aga2p fusion and amyloid β_42_-Aga2p fusion and α-synuclein-Aga2p fusion protein displayed on the cell surface as shown in Figure 1. Additionally, bright-field imaging data showed that there was not any high level of cell aggregation due to NDP interaction (Figure 1B). This is especially important showing that the target proteins, which are prone to aggregate formation in their purified forms, are being displayed in short oligomers and may be as monomeric subunits. Also, fluorescence emission profiles of the cell membranes were also obtained for each of the target proteins based on the immunostaining and intensities. We have verified the display of the target proteins and we have provided with the fluorescence line profiles the presence of the target proteins.

**Figure 1:**
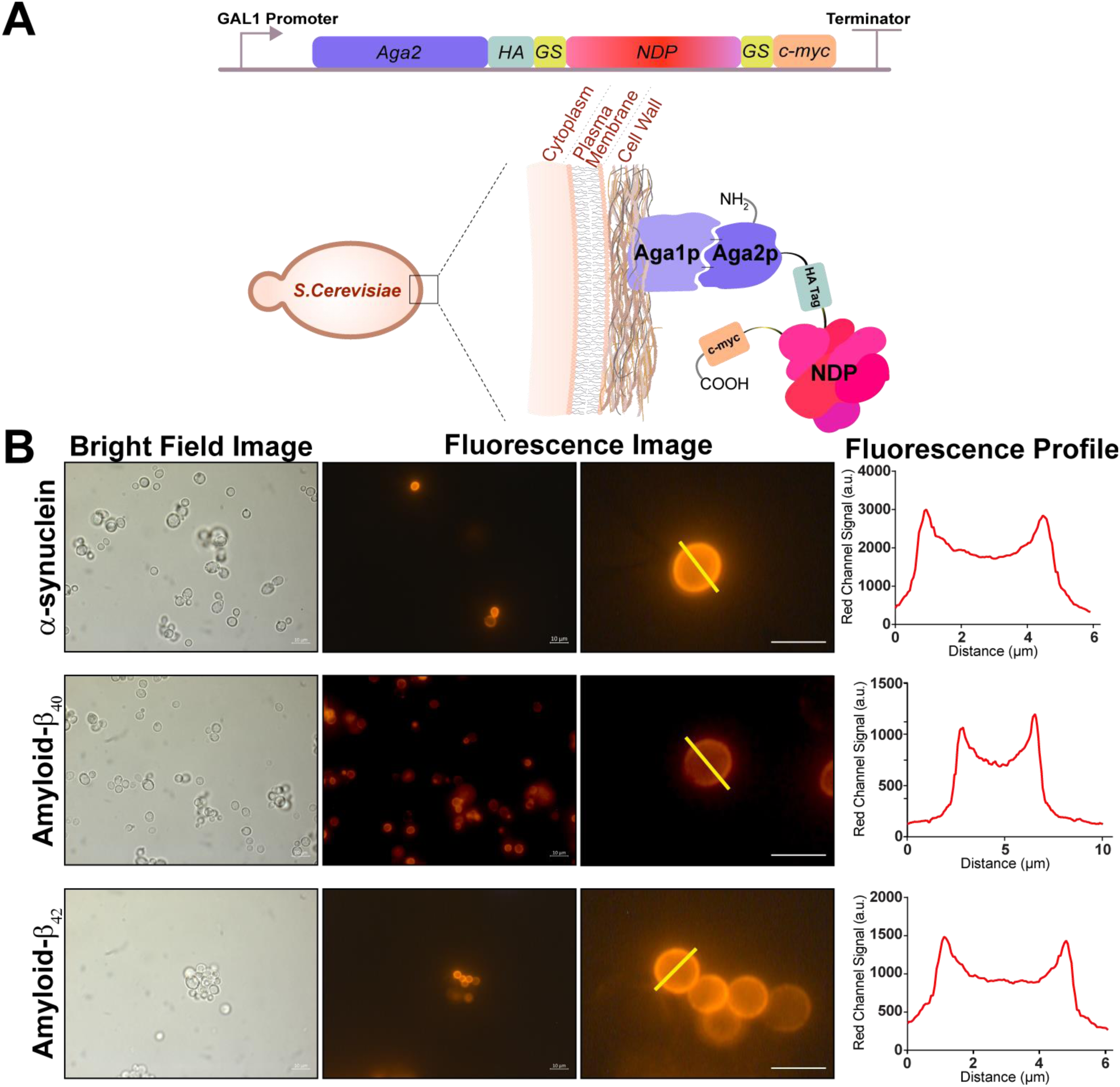
Yeast surface display system was utilized for screening NDPs as monomeric units to select interacting peptides for changing the fibrillization states. (A) Yeast surface display was achieved by fusing NDPs to Aga2p. The expression of NDP-Aga2p occurred after induction by galactose. (B-D) The induced yeast cells expressing NDPs were characterized by ICC method. The immunostaining was applied without any membrane solubilizing agents.

According to the fluorescence emission profile data, each NDP-expressing cell showed homogenized and well-displayed protein expression on the surface, which is used as the verification of the cell surface display of the proteins of interest. It is also evident from the Figure 1, the expressed proteins were transferred to the membrane. Although fluorescence emission profiles were similar throughout the cell membranes, the red fluorescence signal levels for single NDP-expressing cells were different. However, these differences did not effect the use of the final cells as display elements. Furthermore, the variation in expression levels for each NDP in the yeast cell cultures allowed us to asses the efficiency of peptide selection and enrichment ratio of phage suspensions after each cycle of the biopanning.

Both bright field and fluorescence images were taken to analyze the protein expression. The surface profiling was determined from one yeast cells by following the straight yellow line Following the display of the target proteins, we exploited a commercial phage displayed peptide library to select peptides that inhibits aggregation of neurodegenerative amyloids within this study. The commercial library size was around 10^9^ different peptides. After amplification of the library by following the manufacturer’s protocol, the generic panning method for yeast cells was redesigned by using *S. cerevisiae* cells as suspensions instead of immobilization to any surfaces in order to prevent cell loss during iterative washing steps. In our approach, Eppendorf Protein LoBind microcentrifuge tubes (Eppendorf, Hauppauge, NY,USA) were used during biopanning cycles, to prevent unwanted adhesion of proteins and cell on tube walls. Biopanning protocol using a suspension yeast cell display system provided an opportunity to screen a vast number of cells in a relatively larger volume of target cell culture as well as simplicity in the application. Before selecting peptides, preselection was done against the complete empty *S.cerevisiae* cells not displaying any NDPs but the cell mating factors to prevent nonspecific interaction between phage particles and yeast surface displayed elements other than NDPs. Unbound phage particles were eluted and amplified for further screening against NDP-displaying yeast cells. In the -monomeric NDP specific peptide selection step, four iterative cycles of biopanning against each NDP were performed separately (Figure 2A). During each cycle, the input and output phage particles were recorded (Figure 2B). To show the increase in the specificity of phages obtained in the previous cycle, enrichment ratios were calculated by dividing the ratio result (output pfu/input pfu) by the 1^st^ ratio result (Figure 2C). For each biopanning cycle, slight increases were observed in interacted phage particle pool size for α-synuclein and amyloid β_42_ and a sharp increase for amyloid β_40_. These resulted in an increasing trend in enrichment ratios. These results were related to data obtained by ICC. Since there were heterogeneity in induced cell culture. The cultures having more immunostained cells gave a higher enrichment ratio value. Consequently, the increase in amyloid-β_40_ enrichment was higher than amyloid-β_40_ and α-synuclein enrichment ratios. After the last cycle of biopanning was achieved, phage particle elution was tittered to obtain a single clone of each particle. The genomic regions of every phage particle having different peptide sequences in the *pIII* gene of M13 phage encoding for pIII coat protein were sequenced by Sanger Sequencing (Genewiz, NJ, USA). Following sequencing, five of the various peptide sequences were selected for binding α-synuclein and amyloid β_40_. Three additonal peptide sequences were chosen for amyloid β_42_ to prevent peptide-protein interaction and aggregation inhibition (Figure 2D). Peptides were synthesized by solid-state peptide synthesis (Supplementary Methods). Synthesized peptides were dissolved in 1X PBS for further use.

**Figure 2:**
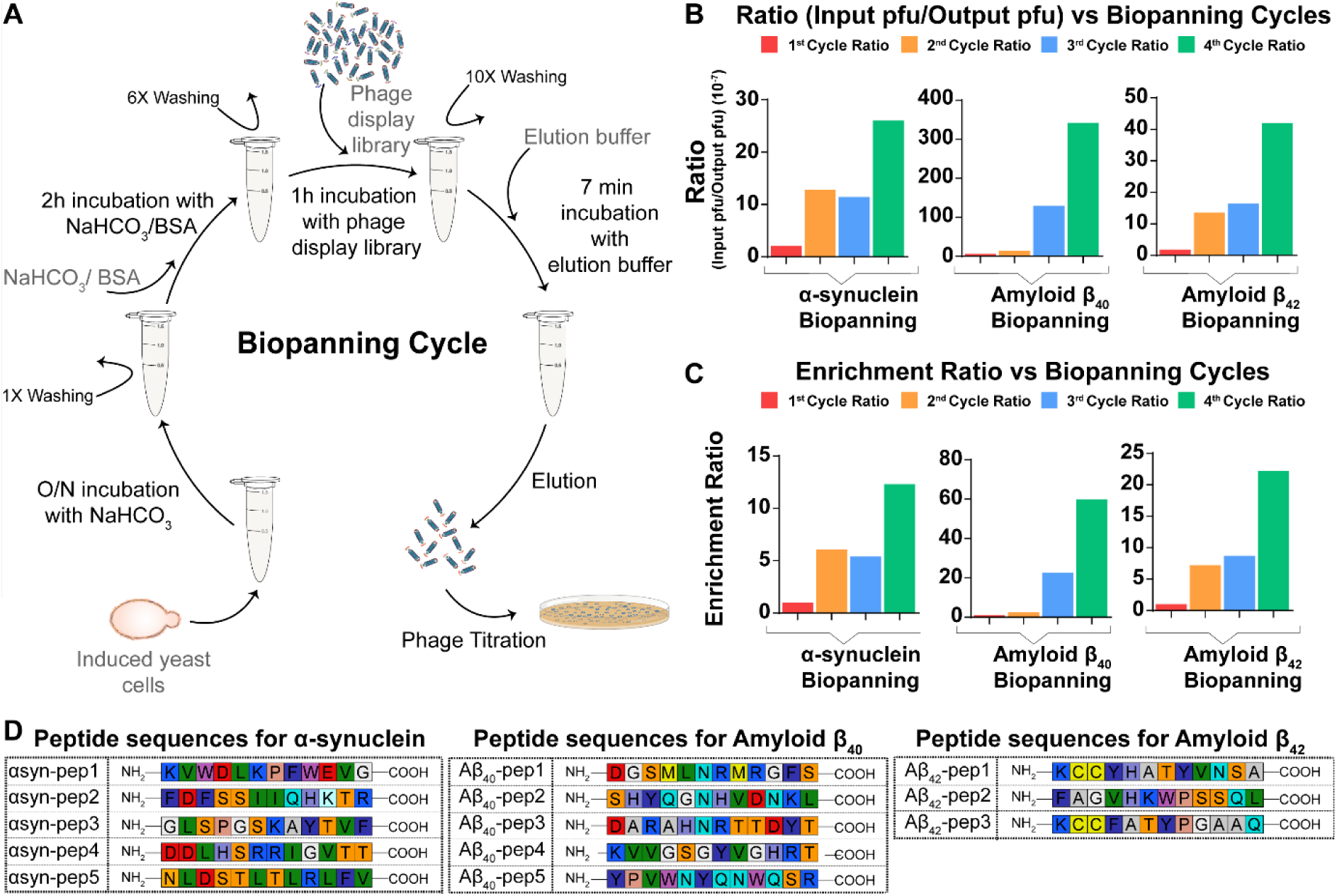
Peptide selection against NDPs was achieved by biopanning protocol with iterative cycles. (A) Yeast surface display-phage display library combination for peptide screening was achieved by optimized biopanning protocol by using yeast cells as suspension and without immobilization. (B) During each cycle, input phage forming units and interacting output phage forming units were recorded to decide the ratio for α-synuclein, amyloid β_40_, and amyloid β_42_ specific phage pool sizes. (C) Enrichment ratios for α-synuclein, amyloid β_40_, and amyloid β_42_ were calculated to show the increase in specificity of obtained phage pool. (D) Selected peptides were sequenced for each NDP. The amino acids were colored according to universal amino acid color codes, which showed their diversity. There were no continuous overlaps for no more than two amino acids.

After completion of the screening and selection of the peptides, we have carried out further analysis of the inhibitory effects. Before characterizing the interaction of peptides with NDPs, monomeric amyloid β_40_ and amyloid β_40_ were used (GenScript) while α-synuclein was purified by fast protein liquid chromatography (FPLC) method after expressed in *E.coli* BL21 (DE3) (Supplementary Figure S2). To increase the solubility, we expressed α-synuclein as a fusion of Glutathione S-transferase (GST). After purification we removed GST tag by TEV protease and obtained monomeric α-synuclein protein for further use (Supplementary Figure S2A-F). The first analysis relied on protein-peptide interaction which was determined by using a mass sensitive technique known as Quartz crystal microbalance device. The QCM-D measures the change of the frequency change of the gold coated quartz crystal sensor upon mass deposition, providing a highly sensitive means of monitoring interactions^40^. In this study we monitored the interaction of inhibitory peptides with NDPs on sensors surface (Figure 3A).

**Figure 3:**
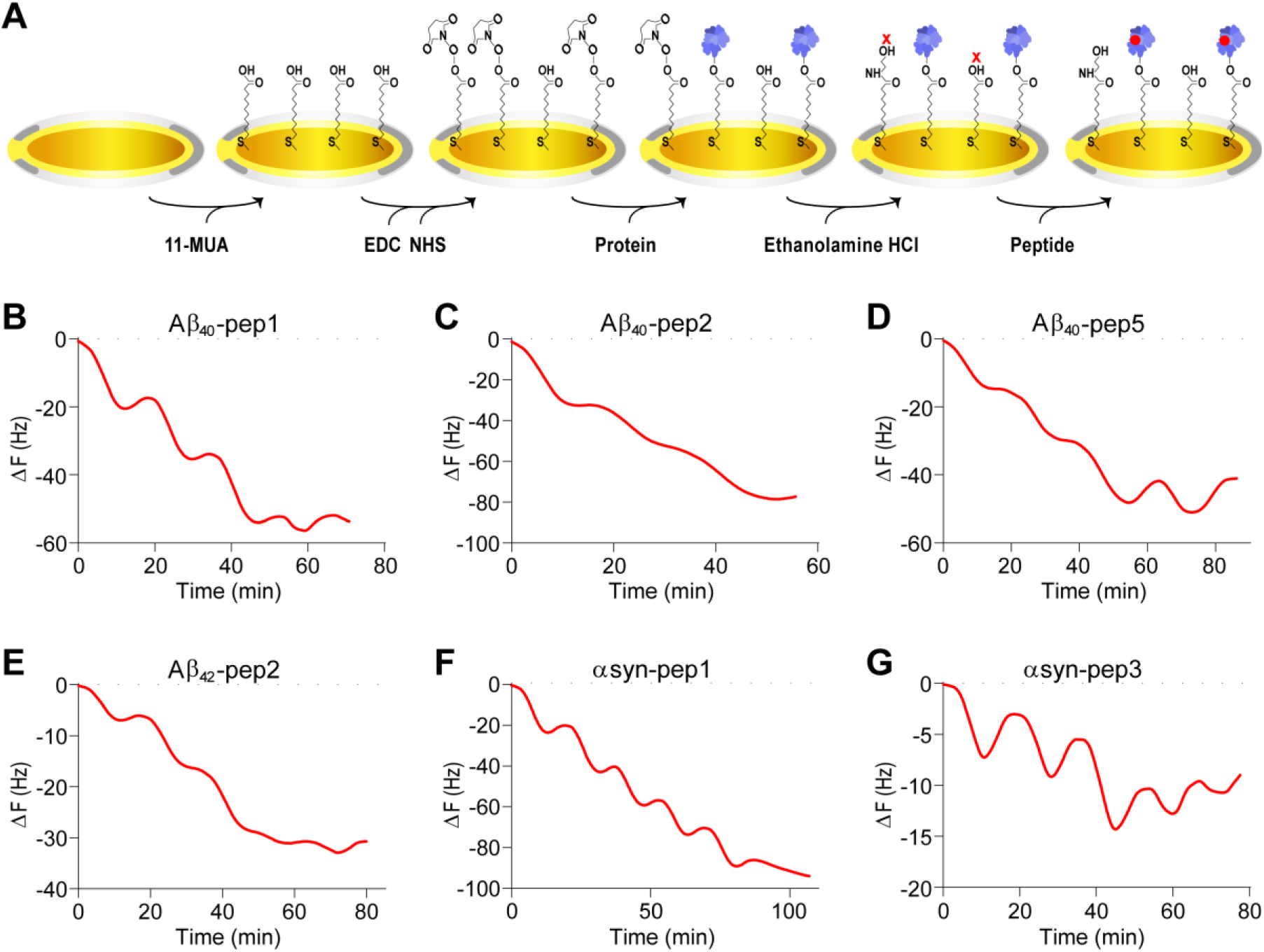
QCM analysis was applied to determine the protein-peptide interaction by evaluating frequency changes. (A) The chip preparation was achieved on a gold chip coated by 11-MUA. EDC/NHS surface activation was applied to immobilize the target protein. The deactivation of the surface was done by 1M ethanolamine HCl addition. Peptides were introduced to the protein-coated chips separately. (B-D) Aβ_40_-pep1, Aβ_40_-pep2, and Aβ_40_-pep5 were delivered onto Amyloid β_40_-coated gold chips, individually. Frequency changes were recorded during peptide additions and washes. (E) Aβ_42_-pep2 was analyzed by delivering onto Aβ_42_-coated chip, and frequency changes were recorded. (F-G) The frequency changes for αsyn-pep1, and αsyn-pep3 with α-synuclein-coated chips were analyzed during each peptide addition and washing.

We analyzed five different amyloid β_40_ peptides and found that each peptide had a unique interaction pattern with amyloid β_40_. Aβ_40_-pep1 showed a well-defined, gradual drop in frequency with each peptide addition, resulting in a mass deposition of 704 ng/cm^2^ onto the amyloid β_40_-coated surface (Figure 3B, Supplementary Figure S3A). For Aβ_40_-pep2, the frequency change decreased continuously between each peptide addition, without a sharp, gradual pattern, resulting in a mass deposition of 786 ng/cm^2^ (Figure 3C, Supplementary Figure S3B).

In the case of Aβ_40_-pep5 addition, there was a gradual drop in frequency for each peptide addition step, and the mass deposition of Aβ_40_-pep5 was calculated as 715 ng/cm^2^ (Figure 3D, Supplementary Figure S3C). For Aβ_40_-pep3, there were gradual but small decreases in frequencies during each peptide addition step, with a mass deposition of 405 ng/cm2 (Supplementary Figure S4A). On the other hand, Aβ_40_-pep4 showed no interaction with the monomeric amyloid β_40_ protein since there was no frequency change between each peptide addition followed by a wash step. The small frequency change corresponded to 9.5 ng/cm^2^ mass deposition, which indicates no significant and specific interaction between Aβ_40_-pep4 and amyloid β_40_ (Supplementary Figure S4B). The small frequency change corresponded to 9.5 ng/cm^2^ mass deposition, which means there was no significant and specific interaction between Aβ_40_-pep4 and amyloid β_40_ (Supplementary Figure S4B).

When we evaluated the interaction of the three peptides with amyloid β_42_, Aβ_42_-pep2 showed a gradual decrease in frequencies between each peptide addition, with a mass deposition of 442 ng/cm^2^ (Figure 3E, Supplementary Figure S3D). Aβ_42_-pep1 showed a more well-defined gradual drop in frequencies with lesser peptide deposition onto the amyloid β_42_ surface, with a value of 134 ng/cm^2^ (Supplementary Figure S4C).

In contrast, there was no optimal pattern for frequency changes for Aβ_42_-pep3 during peptide addition and wash steps. Still, a mass deposition was calculated as 100 ng/cm^2^, although the frequency change during peptide addition was not significant and apparent (Supplementary Figure S4D).

The binding analysis of α-synuclein peptides was conucted with five binding peptides. Among them, αsyn-pep1 showed the most significant gradual drop in frequencies with a mass addition of 878 ng/cm^2^ (Figure 3F, Supplementary Figure S3E). Similarly, αsyn-pep3 interacted with α-synuclein-coated surface with a gradual decrease in frequencies and a mass deposition of 136 ng/cm^2^ (Figure 3G, Supplementary Figure S3E). For the remaining αsyn-peptides, there was no clear and well-patterned gradual change in frequencies, although there were mass depositions of 427 ng/cm^2^, 27.2ng/cm^2^, and 3.8 ng/cm^2^ for αsyn-pep2, αsyn-pep4 and αsyn-pep5, respectively (Supplementary Figure S4E-G). Thus, peptides that had an affinity to NDPs caused a decrease in frequency for each peptide addition step, with a considerable mass deposition onto NDP-coated surfaces. Additionally, we demonstrated that the well-interacted peptides with the NDP-coated surfaces exhibited clear changes in frequencies.

### Imaging of the inhibitory effect of the peptides on Amyloid Formation

The evaluation of peptide interactions and their effects on the fibrillization of NDPs was carried out using AFM topographical analysis. The analysis involved the evaluation of up to seventy individual fibrils using the FiberApp analysis tool. Each distinct fibril or fibril branch (spikes) was evaluated and compared with itself only because the background signals were high in each data due to high and nonuniform aggregation and fibrillization^41^. The AFM determined more coherent heights, but the comparison for heights and contour lengths was determined by FiberApp. The results of the analysis are presented in Figure 4 and Supplementary Figure S5.

**Figure 4:**
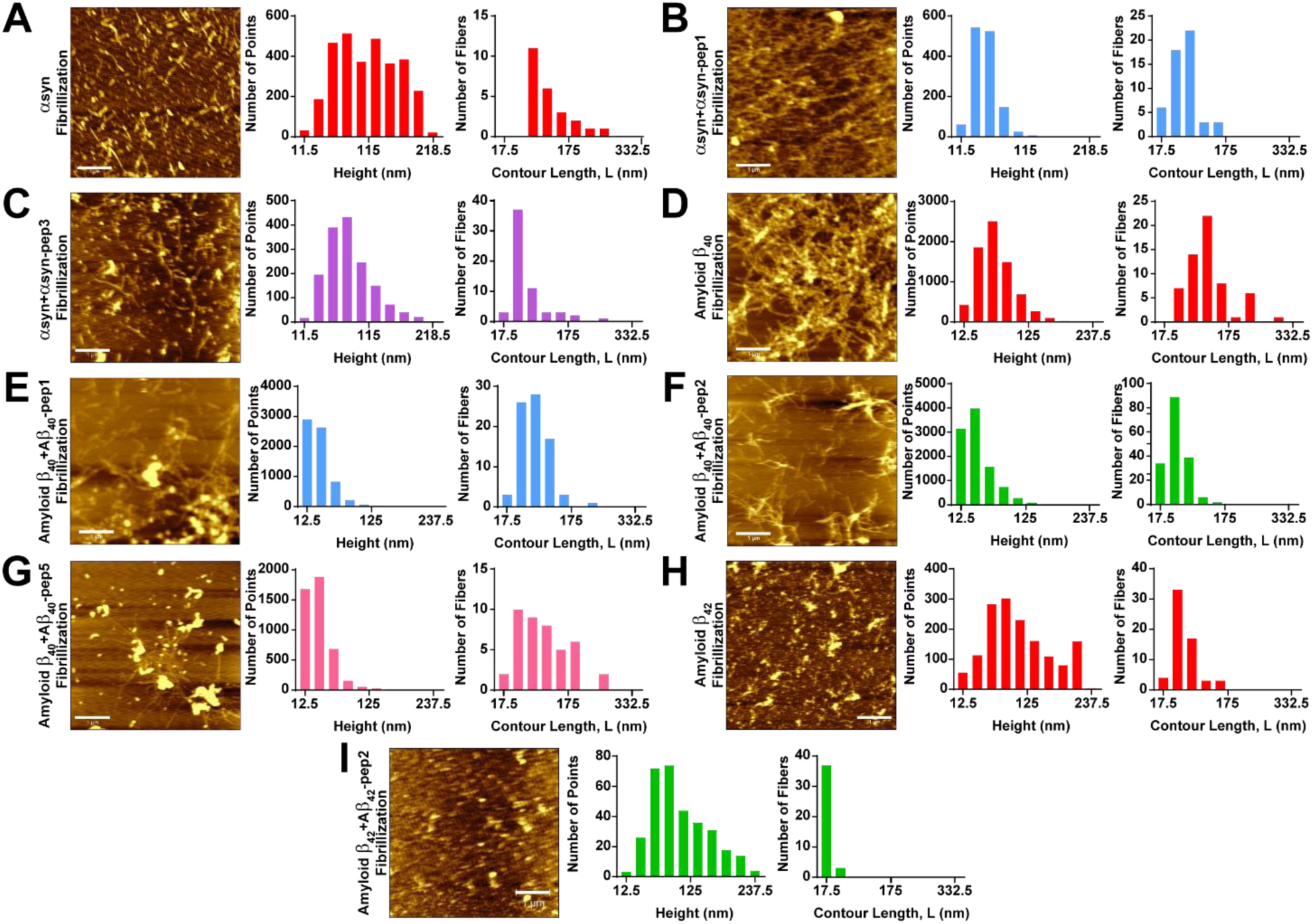
Fibrillization states of monomeric NDPs with and without peptides were analyzed by AFM in tapping mode under dry conditions to detect fibrillization states. (A) α-synuclein fibrillization was achieved without any peptides. The heights and lengths were analyzed from detectable fibrils. (B-C) α-synuclein with αsyn-pep1, and αsyn-pep3 fibrillization assay was analyzed, and heights and lengths from distinct fibrils were determined individually. (D) Amyloid β_40_ fibrillization product without any peptide addition was analyzed, and the heights and lengths of the detectable fibrils were determined. (E-G) Amyloid β_40_ fibrillization products with Aβ_40_-pep1, Aβ_40_-pep2, and Aβ_40_-pep5 were analyzed, and heights and lengths were determined, respectively. (H) Amyloid β_42_ fibrillization was evaluated. The heights and lengths were decided from distinctively observed spikes. (I) Amyloid β_42_ fibrillization products with Aβ_40_-pep2 was visualized by AFM as seeds whose heights and lengths were determined by FiberApp.

The analysis of α-synuclein fibrillization revealed various profiles for both α-synuclein fibrils alone and in mixtures with α-synuclein inhibitory peptides. AFM images of α-synuclein fibrillization showed the presence of individual seeds and non-identical fibril branches of varying sizes (Figure 4A). The seeds and fibril spikes were distinguishable and easily countable, and their intensities were relatively high, indicating large volume sizes. The height and contour length (L) of all detectable α-synuclein fibrils were analyzed, and there was significant heterogeneity in both parameters due to the state of fibrillar maturation. However, larger and longer fibrils were considered mature α-synuclein fibrils.

Comparing α-synuclein fibril AFM data with α-synuclein+αsyn-pep1 fibrillization, the latter occurred in a crowded mesh-like structure with lower intensities and shorter calculated heights and detectable individual fibril spike lengths (Figure 4B). Despite this, the mesh-like fibril structures made it difficult to detect separate and individual fibrils, indicating αsyn-pep1’s affinity for α-synuclein by increasing fibrillization. Conversely, AFM image data for α-synuclein+αsyn-pep3 fibrillization showed a positive inhibitory effect on fibrillization, with a relatively higher seed amount and fewer fibril branches (Figure 4C). Although the lengths of the fibrils were diverse, the contour lengths were generally short, indicating αsyn-pep3’s inhibitory effect on fibrillization.

When the fibrillization of α-synuclein with αsyn-pep2, αsyn-pep4, and αsyn-pep5 was analyzed individually, AFM image data showed mesh-like fibrillization with diverse fibril intensities and short detectable individual fibril lengths (Supplementary Figure S5A-C). However, the main fibrils creating the aggregation mesh would be long but not distinguishable as single fibrils. Thus, these three peptides were also considered interacting peptides with positive effects on fibrillization. End-point Thioflavin T (ThT) fluorescence measurements were also done to support this evaluation, where αsyn-pep5 had a higher impact on fibrillization with higher ThT fluorescence than α-synuclein fibrillization ThT fluorescence, while αsyn-pep3 had the lowest ThT fluorescence than monomeric α-synuclein (Supplementary Figure S6A).

In the case of amyloid β_40_, AFM fibrillization data for amyloid β_40_ and amyloid β_42_ separately showed different fibrillization profiles. Amyloid β_40_ alone exhibited proper fibrillization profiles with mesh-like fibrils having seeds in the fibril branch centers (Figure 4D). Although detecting individual fibrils was challenging, the selected fibrils were relatively higher in height and fibril length as expected. For the fibrillization of amyloid β_40_ + Aβ_40_-pep1, there were short fibril spikes with diverse-sized amyloid aggregates (Figure 4E). According to the fibril intensities, the aggregates were relatively less intense, and the fibril sizes possessed considerably shorter lengths with less thickness than amyloid β_40_ fibrils. The amyloid β_40_ + Aβ_40_-pep2 AFM data also showed decreased fibrillization with shorter and less amount of fibril spikes (Figure 4F). Also, seed aggregates were observed in fewer amounts over the whole fibrillar structures. Although detectable branches were high in amount, most possessed small contour lengths, and heights were prone to be small. When the fibrillization of amyloid β_40_ + Aβ_40_-pep5 was analyzed, seeds dominated the fibrillization profile (Figure 4G). Fibril branches were highly separate and individually countable, and long as observed in amyloid β40 fibrils, although the amount was low. The contour lengths were diverse, and heights tended to be less in amount. The overall profile of amyloid β_40_ + Aβ_40_-pep5 mix did not fit or show complete similarities with either fibrillization profiles or inhibited fibrillization. The existence of intense and large volumes of seeds and long and separate fibrils exhibited more atypical fibrillization than the others. For amyloid β_40_ + Aβ_40_-pep3 and amyloid β_40_ + Aβ_40_-pep4 fibrillization AFM image data, fibrils obtained in the existence of Aβ_40_-pep3 were less crowded than with the existence of Aβ_40_-pep4 (Supplementary Figure S5D-E). However, according to the data, Aβ_40_-pep3 and Aβ_40_-pep4 had less positive effects on the inhibition of aggregation and fibrillization. Further, ThT fluorescence emission results showed that amyloid β_40_ + Aβ_40_-pep5 gave a higher signal than amyloid β_40_ fibril alone and with the peptide mixes. However, amyloid β_40_ + Aβ_40_-pep1 and amyloid β_40_ + Aβ_40_-pep2 fibril samples gave higher signals than amyloid β_40_ + Aβ_40_-pep3 and amyloid β_40_ + Aβ_40_-pep4 mixes as well as monomeric amyloid β_40_. The reason for this might be originated from the blockage of cross β structures, where ThT binds, by peptides (Supplementary Figure S6B).

AFM analysis of amyloid β_42_ for fibrillization inhibition showed the effect on the fibrillization growth of the peptides (Figure 4H). Unlike amyloid β_40_ fibrils, amyloid β_42_ fibrils produced fewer mesh-like fibrillar structures. The seeds were in high amounts, which made them the most dominant structure in AFM analysis. Spikes originating from seeds were observed in considerable amounts. Also, there were also individual fibrils, although the lengths of the fibrils were shorter than those observed in cases of amyloid β_40_ and α-synuclein. Nevertheless, fibril growth was started and showed immature amyloid fibril structures. The fibrillization done with amyloid β_42_ and Aβ_42_-pep1 showed fewer seeds and no fibrillar structure (Supplementary Figure S5F). Seeds had no branching or spike-like structures. The seed sizes were diverse, with higher heights and short aggregate lengths.

AFM analysis showed that fibrillization inhibition of amyloid β_42_ had an effect on the growth of peptide fibrils (Figure 4H). In contrast to amyloid β_40_ fibrils, amyloid β_42_ fibrils produced fewer mesh-like fibrillar structures, with the dominant structure being seeds in high amounts. Considerable spikes originating from seeds were also observed, along with individual fibrils, though the lengths of the fibrils were shorter than those observed in cases of amyloid β_40_ and α-synuclein. Nevertheless, fibril growth was initiated and showed immature amyloid fibril structures. Fibrillization of amyloid β_42_ with Aβ_42_-pep1 resulted in fewer seeds and no fibrillar structure (Supplementary Figure S5F). Seeds had no branching or spike-like structures, and their sizes were diverse, with higher heights and short aggregate lengths.

Even though the length of the amyloid β_42_ units had similar values in the products of the amyloid β_42_+Aβ_42_-pep2 mixture, some of the seed structures were prone to branching. The height distribution was in a wide range for the detectable spike-like structures. To determine fibril formation depending on β-sheet structures, ThT fluorescence emissions were measured. Fibrils produced only from amyloid β_42_ emitted more fluorescence than the rest, followed by amyloid β_42_+Aβ_42_-pep1. Monomers of amyloid β_42_ produced more ThT fluorescence signal than amyloid β_42_+Aβ_42_-pep3 and amyloid β_42_+Aβ_42_-pep2, respectively (Supplementary Figure S6C). The higher signal from amyloid β_42_+Aβ_42_-pep3 was expected, and it was predicted that amyloid β_42_+Aβ_42_-pep1 would possess a lower ThT signal relatively than the other amyloid β_42_+Aβ_42_-pep mixtures.

The reason for the unexpected results can be attributed to several factors. Firstly, Aβ_42_-pep3 may have blocked the β-sheets that ThT interacts with, resulting in lower ThT fluorescence signals. Although fibrils were observed in the amyloid β_42_+Aβ_42_-pep3 samples, the presence of Aβ_42_-pep3 may have interfered with the ThT detection. Additionally, the QCM results indicated that the affinity of Aβ_42_-pep3 to amyloid β_42_ was low, which may have contributed to the reduced ThT signal. Secondly, Aβ_42_-pep1 may have intrinsic fluorescence, leading to higher fluorescence signals despite the absence of fibrillization observed in AFM analysis. Although the deposition mass amount of Aβ_42_-pep1 was low according to QCM results, it may still be useful in designing peptide-based drugs or sensor molecules for detection. Finally, in the case of amyloid β_42_+Aβ_42_-pep2, the ThT signal was the lowest, possibly due to the fact that Aβ_42_-pep2 had a higher affinity to amyloid β_42_, as confirmed by the AFM image showing no distinct fibrils.

The biopanning protocol utilized in our study involved the use of yeast cells displaying neurodegenerative proteins as monomers. This approach allowed for the proper folding of NDP monomers on the surface, along with post-translational modifications, and enabled the selection of peptides based on their expression levels, which increased during the process. Following the selection of phage particles from the library, we obtained peptide sequences, from which we selected five peptides for α-synuclein and amyloid β_40_, as well as three peptides for amyloid β_42_. We then analyzed these peptides using QCM and AFM to evaluate their interaction efficiency and their effect on fibrillization, respectively.

The QCM analysis with the same concentration of peptides showed that some of them showed a gradual decreasing pattern for ΔF value with diverse mass depositions. αsyn-pep1, αsyn-pep3, Aβ_40_-pep1, Aβ_40_-pep2, Aβ_40_-pep5, and Aβ_42_-pep2 showed high and more gradual mass deposition than the other peptides. Based on these results, fibrillization production was completed for all monomeric units with and without peptides. The fibrillization products showed the amyloid oligomerization and fibrillization states and the effect of peptides on the fibrillization event. αsyn-pep3, Aβ_40_-pep1, Aβ_40_-pep2, Aβ_40_-pep5, Aβ_42_-pep1, and Aβ_42_-pep2 had positive effect on fibrillization. However, ThT results for Aβ_40_-pep1, Aβ_40_-pep2, and Aβ_42_-pep1 conflicted with the fibrillization analysis by AFM. Still, each of the peptides had an effect on fibril formation by promoting or inhibiting the process. Briefly, αsyn-pep3, Aβ_40_-pep1, Aβ_40_-pep2, and Aβ_42_-pep2 were good candidates for peptide-based drug developments for blocking fibrillization from the early stages of the disease conditions.

Further, αsyn-pep1, αsyn-pep2, αsyn-pep4, αsyn-pep5, Aβ_40_-pep3, Aβ_40_-pep4, Aβ_40_-pep5, Aβ_42_-pep1, and Aβ_42_-pep3 can be engineered and utilized for sensor molecules for detecting the disease conditions. For the detection, the over-expression of amyloid units can interact more with the sensor candidate peptides and give higher intense fibril formation as well as ThT fluorescence signals.

Overall, the peptides selected and analyzed in our study showed a significant impact on amyloid fibril formation, highlighting the potential of our yeast surface display and phage display library approach for discovering peptide-based therapeutics and molecular diagnosis tools for neurodegenerative diseases. The simple application protocol, high yield of displayed proteins, and short application time of this combination make it a promising approach for developing new drug molecules or detection platforms. Since early diagnosis and halting disease progression are crucial in treating Parkinson’s and Alzheimer’s diseases, the use of surface display platforms to select novel peptide molecules holds great promise in diagnosis and blocking these diseases.

## Methods

### Materials

For surface display, The *Saccharomyces cerevisiae* strain EBY100 (ATCC® MYA-4941™) (a GAL1-AGA1::URA3 ura3-52 trp1 leu2Δ1 his3Δ200 pep4::HIS2 prb1Δ1.6R can1GAL) was purchased. Ph.D.™-12 Phage Display Peptide Library Kit (New England BioLabsⓇ Inc., MA, USA) was preferred in order to select aggregation inhibitory peptide selection. For the in vitro characterization studies, amyloid β_40_ and amyloid β_42_ were purchased (Genscript, Cat.#RP10004 and #RP10017, respectively). For the interaction studies, QCM-D gold chips were purchased from Biolin Scientific (QSX 301).

### Construction of plasmids

Yeast surface display construct was done by using pETcon (-) plasmid, which was a gift from Andrew Scharenberg (Addgene plasmid # 41522 ; http://n2t.net/addgene:41522 ; RRID:Addgene_41522). Amyloid β_40_ and amyloid β_42_ genes were extended and amplified by overlap extension PCR (supplementary figure), and cloned by the Gibson assembly method in the pETcon (-) plasmid. α-synuclein gene was synthesized as a fragment gene by Genewiz (Genewiz, NJ, USA) and cloned in pETcon (-) plasmid by the Gibson assembly method. sfGFP genes were amplified by PCR from pJT119b plasmid which was kindly gifted by by Jeffrey Tabor (Addgene plasmid #50551), and cloned into pETcon-alpha syn, pETcon-amyloid β_40_ and pETcon-amyloid β_42_ by the Gibson assembly method. For protein expression and purification in *E*.*coli*, α-synuclein was amplified by PCR. As the backbone plasmid, pET22b-6h-GST-TEVp was digested with BamHI and XhoI restriction enzymes in order to remove TEVp gene (Addgene plasmid #172887). α-synuclein was cloned into pET22b-6h-GST backbone by Gibson assembly method. All cloned constructs were chemically transformed into *E.coli* DH5α PRO strain. All constructs were verified by Sanger sequencing (Genewiz, NJ, USA).

### Transformation of EBY100 cells

The *Saccharomyces cerevisiae* strain EBY100 (ATCC® MYA-4941™) (a GAL1-AGA1::URA3 ura3-52 trp1 leu2Δ1 his3Δ200 pep4::HIS2 prb1Δ1.6R can1GAL), Trp-Leu-was utilized for the surface expression of α-synuclein, amyloid β_40_, and amyloid β_42_. pETcon constructs were transformed into the EBY100 cells via electroporation. Electrocompetent EBY100 cells were prepared following the protocol developed by Suga and Hatakeyama. Briefly, EBY100 cells were grown overnight to reach a cell density value of around 1×10^7^ cells/ml. Cell culture was placed on ice for 15 minutes and centrifuged at 4000 g for 5 minutes. Pellet was washed three times in cold sterile double-distilled water. After washing steps, cells were resuspended in ice-cold freezing buffer (2M sorbitol, 10 mM CaCl2 and 10 mM HEPES (pH 7.5)) as the cell concentration became 5×10^8^ cells/ml. 100 µl aliquots of cell suspension were transferred into the 1.5 ml sterile cryotubes. After thawing electrocompetent cells in a 30°C water bath, 1 ml of 1 M sorbitol was added. Cells were centrifuged and the pellet was dissolved in 1 M sorbitol to adjust cell density to 1-2×10^8^ cells/ml. After the addition of purified plasmid, cell suspension was transferred into a cuvette with a 2 mm gap size. Following the electrical pulse (voltage: 2 kV, resistance: 200 Ω, capacitance: 25 µF), 1 ml of ice-cold sterile 1 M sorbitol was immediately added onto the cells. 100-200 µl of the cell mixture was spread onto the synthetic-dropout (SD)-agar plates to be incubated at 30°C for 2-3 days.

### Induction of yeast cells for surface display

In order to induce surface display of the neurodegenerative disease related proteins, EBY100 cells were first grown in 3-5 ml of synthetic-drop out media without tryptophan including 2% dextrose (SD-CAA) with constant shaking at 260 RPM at 30°C. OD_660_ value was measured and cells were transferred into fresh SD-CAA and the cell density was arranged to be approximately 4.6×10^6^ cells/ml. The concentration was sufficient to reach OD_660_ value to 1 in four-hour incubation at 260 RPM at 30°C. After incubation, cell suspension was centrifuged at 4000 g for 5 minutes and cell pellet was dissolved in selective induction medium (1.92 g/L Synthetic Drop-out medium without tryptophan, 6.7 g/L Yeast Nitrogen Base without amino acids, 2% galactose, 0.2% glucose, 1X phosphate buffer, and 100 µg/ml ampicillin). Cells within the synthetic drop-out induction medium with galactose (SD-GAA) were incubated at 260 RPM at 20°C for 16-20 hours.

### Immunostaining of yeast cells

1 × 10^6^ of induced yeast cells were centrifuged and induction medium was discarded. After washing the cell pellet with 1X PBS, cells were resuspended in 1 mL of blocking solution (1% BSA in 1X PBS [w/v]). For blocking, the cells were incubated at room temperature for 2 hours. At the end of the blocking incubation, cells were centrifuged and blocking solution was discarded. Then, cells were washed three times with 1X PBS. After washing, cells were resuspended in 250 μL1% BSA in 1X PBS containing Myc-tag Mouse monoclonal antibody (Cell Signaling, 9B11, Cat.#2276) with a ratio of 1:500. Primary antibody incubation was done at room temperature for 3 hours. Following the primary antibody incubation, cells were washed with 1X PBS three times. Then, cells were resuspended with 250 μL1% BSA in 1X PBS containing 1:500 Goat Anti-Ms IgG (H+L) Cross Adsorbed Secondary Antibody, DyLight550 conjugate (Thermo Scientific, SA5-10173). The secondary antibody incubation was done at room temperature for 3 hours. At the end of the incubation, cells were washed with 1X PBS three times, and resuspended in 250 μL of 1X PBS. The labelled cells were placed onto positively charged slides and examined with Zeiss Axio Scope.A1 microscope

### Biopanning against yeast expressing neurodegenerative proteins on the surface

To select aggregation inhibitor peptides, yeast surfaces displaying neurodegenerative proteins were produced as explained in the section on EBY100 induction. Ph.D.™-12 Phage Display Peptide Library Kit (New England BioLabs® Inc., MA, USA) was amplified as following the recommended protocols by the manufacturer. Before selecting ligand peptides, the negative selection was done by using Aga2p displaying EBY100 cells. Unbound phages were eluted and amplified as specified in the manual of the library. The unbound phage library was used to select peptides against EBY100 cells displaying neurodegenerative proteins. For all biopanning methods (for both negative selection and the selection), 10^11^ yeast cells were used at the beginning of each biopanning cycle. All centrifugation steps for yeast cells were done at 3000 g for 3 minutes. The number of phage particles used in each cycle of biopanning was 10^9^. Except for the negative selection, bound phages were eluted and amplified for further biopanning cycles. For each biopanning cycle was done by using Protein LoBind® Tubes (Eppendorf) in order to avoid nonspecific interactions of proteins with the tubes during biopanning. At the beginning of each cycle, 10^11^ yeast cells were centrifuged down and the pellet was washed with 1 mL 0.1 M NaHCO_3_, pH 8.6 coating buffer. After centrifugation, the pellet was again resuspended with 1,5 mL of the same buffer and incubated at +4°C overnight onto the rotator. The next day, cells were centrifuged and the coating buffer was removed. The cell pellet was washed with 1,5 mL 1X TBS-T and centrifuged. Then, the pellet was resuspended with 1,5 mL 0.1 M NaHCO_3_, pH 8.6 containing 5 mg/mL BSA, and incubated for 2-4 hours at 4°C on rotator. At the end of the blocking, cells were centrifuged and washed 6 times with 1X TBS-T buffer. When wash steps were finished, 10^9^ phage particle was added onto the cell pellet and the volume was completed to 1 mL with 1X TBS. For achieving interaction, cells and phages were incubated at room temperature for 1 hour on rotator for mixing them thoroughly. At the end of phage binding incubation, cells were centrifuged and washed 10 times with 1.5 mL 1X TBS-T to get rid of the unbound phages. For elution of the unbound phages, 1 mL 0.2 M Glycine-HCl, pH 2.2 elution buffer was added to cells and incubated at room temperature for 7 minutes with gentle shaking. To get the phage elution, the supernatant was transferred to a new tube and 150 μL neutralization buffer (1M Tris-HCl, pH 9.1) was added.

### Phage titering

Overnight culture of *E.coli* ER2738 cells was diluted with a ratio of 1:100 into LB containing tetracycline antibiotic. At the time of OD_600_ value reached 0.5, 200 μ Lof cells were transferred into sterile tubes. Meanwhile, amplified phages and eluted phages were diluted with a dilution factor of 10^−8^ to 10^−12^ and 10^−2^ to 10^−4^, respectively. For each dilution, 10 μL samples were taken and added to 200 μL of cells. Phages were incubated with cells for 1-5 minutes at room temperature in order to infection occurred. At the end of the incubation, cell-phage samples were mixed with 45°C, 3 mL top agar. Top agar containing phage-cell mixture, then, spread onto LB agar plates containing X-Gal, IPTG, and tetracycline. The plates were incubated at 37°C overnight to complete blue/white screening. The next day, the number of plaques was counted to calculate the enrichment ratio.

### The eluted phage and single phage plaque amplification

To use eluted phages for the next round of biopanning, overnight culture of *E.coli* ER2738 was diluted with a ratio of 1:100 into 20 mL LB containing tetracycline. The diluted culture was incubated until OD_600_ reached 0.01-0.05. After getting the appropriate OD_600_ value, eluted phages were added onto the culture, and incubation was done at 37°C, 200 rpm for 4.5 hours to propagate phage elution. Then, the culture was centrifuged at 12000 rpm, 4°C for 10 minutes. The supernatant having amplified phages was transferred into a new tube 5 mL of 20% PEG-8000/ 2.5 M NaCl solution was added to phage precipitation after overnight incubation at 4°C. Next, phages were centrifuged at 12000 rpm 4°C for 15 minutes. The supernatant was discarded and the finger-like pellet was resuspended with 1 mL 1X TBS. Additional phage precipitation was applied by the addition of 200 μL 20% PEG-8000/ 2.5 M NaCl solution. The phage suspension was incubated on ice for 1 hour. At the end of the incubation, centrifugation was done at 14000 rpm and 4°C for 5 minutes. The phage tittering was done for this amplified phage solution. For using a single phage clone for further experiments, the single phage plaque was selected with a tip and inoculated to 1 ml of *E.coli* ER2738 culture at OD_600_ value between 0.01-0.05, which was prepared with overnight culture as diluted with a ratio of 1:100 into 20 mL LB containing tetracycline. The amplification in the 1 mL culture was carried out at 37°C for 4.5 hours at 200 rpm. Then, the cell culture was centrifuged at 14000 rpm for 30 seconds. The upper 800 μL of supernatant was taken gently without disrupting the cell pellet. 20 μL from 800 μL single phage suspension was further amplified by following the same protocol for eluted phage amplification. All phage suspensions were stored at 20°C with 50% glycerol (1:1 V/V) for long term.

### Expression and purification of α-synuclein

For protein purification, pET22b-α-synuclein-TEV-GST-6H plasmid was transformed chemically into *E.coli* BL21 (DE3) strain. Single colony for α-synuclein expression was picked and inoculated into 1L of autoinduction media^42^. Overnight grown and induced cells having pET22b-α-synuclein-TEV-GST-6H plasmid was harvested for purification steps. Cells were centrifuged at 8000g for 5 min. Next, pellet was resuspended in 100 mL lysis buffer (20mM imidazole, 500 mM NaCl, 20 mM sodium phosphate, pH 7.4). PMSF was added to cell suspension. To lyse the cells, first liquid nitrogen freezing and thawing were applied five times. Then sonication was done by applying 800-900 kJ energy with 30% power for about 2 minutes of 10 s on/10 s off. After lysis of the cells, centrifugation was done at 15000g for 30 minutes. The lysate was transferred to a fresh tube after being filtered with 0.45 um syringe cellulose filters. The protein purification was achieved by using the HisTrap nickel column (GE Life Sciences 17524701) for FPLC (ÄKTA start protein purification system) as the manufacturers’ specified protocol. At the end of the protocol, the elutions having alpha syn-GST protein were collected (Figure S2)

### Producing monomers of α-synuclein

To get the monomeric version of α-synuclein, purified alpha-syn-TEV Recognition Site-GST-6His protein was transferred to the TEV reaction buffer (50 mM Tris-HCl [pH 8.0], 0.5 mM EDTA, and 1 mM DTT). TEV protease reaction was done by using 500 ug/ml alpha syn, 100 ug/mL homemade TEV protease-GST-6His, 0.02% Sodium Azide, 0.1 mM DTT at room temperature overnight. The monomeric proteins were then purified by using the HisTrap nickel column (GE Life Sciences 17524701) for FPLC (ÄKTA start protein purification system) as the manufacturers’ specified protocol. The unbound proteins were collected since α-synuclein had no tag to captured by the HisTag column except TEV Protease-GST-6His and GST-6His from alpha-syn-TEV Recognition Site-GST-6His. The monomeric α-synuclein was transferred to 1X PBS or phosphate buffer for QCM-D analysis or fibrillization assay, respectively.

### Fibrillization assay for amyloid β_42_, amyloid β_42_ and α-synuclein

To produce seeds and check the interaction of peptide-protein, 100 μM monomeric α-synuclein in phosphate buffer, pH 6 was incubated with and without 10 μM peptides separately in Grant PCMT Microtube/microplate Thermo-Shaker at 40°C with 850 rpm for 72 hours. For further producing fibrils of α-synuclein, 300 μM monomers in phosphate buffer, pH 6 were incubated with 100 μM seeds and the mix was incubated at 40°C with 850 rpm for 72 hours.

For the fibrillization of amyloid β_40_ in 0.1M sodium bicarbonate buffer, pH 9 and amyloid β_42_ in PBS, pH 7.4,100 μM protein, 100 μM proteins were added into PBS containing 2% DMSO. To increase the fibrillization, HCl was added to catch the 20 mM final concentration of HCL. The fibrillization was achieved after 24 hours incubation at 37°C. Monomers were also incubated with 250 μM peptides separately

### SDS-PAGE and Western blotting

For SDS-PAGE and western blot analysis, protein samples were heated at 95°C with 1X SDS loading dye. The gel electrophoresis was done with 15% SDS-polyacrylamide gel. After the run was completed, the gel was stained with Coomassie blue staining solution by heated with a microwave oven, then shaken for 5 minutes at room temperature. The stained gel was then placed in the destaining solution (60% ddH_2_O, 30% methanol, and 10% acetic acid) until the excessive dye was removed and bands were visible. For western blot analysis, the samples in the gel were transferred to the PDVF membrane by Trans-Blot Turbo (Bio-Rad) system. The membrane was blocked with 5% milk in TBS-T for 1 h at room temperature. After blocking, the membrane was incubated in 5% milk in TBS-T containing 1:5,000 primary antibodies for 1 hour at room temperature. The membrane was washed with 1X TBS-T and incubated with horseradish peroxidase (HRP)-conjugated secondary antibody for 1 hour at room temperature. At the end of incubation, the membrane was washed with 1X TBS-T. Incubation with ECL substrates (Bio-Rad 170-5060) was done to visualize the membrane. The images were taken by Vilber Fusion Solo S. For α-synuclein oligomers, Anti-α-synuclein Antibody, oligomer-specific Syn33 (Cat.# ABN2265, Millipore) and Goat Anti-Mouse IgG H&L (HRP) (ab6789, Abcam) were used as primary and secondary antibodies, respectively. For HisTag recognition, 6x-His Tag Monoclonal Antibody (HIS.H8) (MA1-21315, Thermo Scientific) and HRP-conjugated goat anti-mouse secondary antibody (Abcam ab6789-1 MG) were used as primary and secondary antibodies, respectively.

### QCM analysis

The QCM gold sensor (Biolin Scientific QSense QSX 301 Gold) surface was cleaned, immersed in 20 mM 11-mercaptoundecanoic acid (11-MUA), and activated by EDC/NHS coupling reaction followed by protein immobilization, as previously described [recep]. For each QCM analysis of amyloid β proteins and α-synuclein, 50 ug and 500 ug proteins were introduced to the chamber, respectively. After the deactivation of the surface was done, 1000 μM of one type of peptide at once was introduced to the chamber sequentially. Between each peptide addition to the chip chamber, the chip was washed with 1X PBS. First, third, fifth, seventh, and ninth overtones for frequency values were recorded during the run. Each plot was obtained by depending on the average of ΔFs from third, fifth, seventh, and ninth overtones. The mass depositions were calculated by the formula of Δm=-C(ΔF/n), where Δm is the change in the mass (ng.cm^−2^), C is the mass sensitivity constant that depends on the chip specification (17.7 ng.cm^−2^ for QSX 301 gold QCM chip), ΔF is the change in the resonance frequency (Hz), and n is the number of harmonics. Mass deposition values were obtained according to the data from fifth overtone.

### AFM analysis

5 μLof fibrils, fibrils with peptides, and monomers were added to the silica wafer, and 200 μL of 2.5% glutaraldehyde was added to each wafer. After overnight incubation of wafers at +4 C, wafers were washed with 1X PBS and then ddH_2_O for 5 minutes in a shaker at room temperature for three times. Then wafers were washed sequentially with 25%, 50%, and 75% ethanol (EtOH) for 5 minutes in a shaker at room temperature. Finally, 100% EtOH washing was done for 5 minutes in a shaker at room temperature three times. After washing, wafers were first air-dried, and argon-gas dried before measuring. AFM images were obtained by MFP-3D Origin AFM (Oxford instruments). Diamond-like carbon (DLC) coated AFM cantilever tip having a spring constant of 40 N/m with resonance frequency of 300 kHz was preferred. Surface topography of the samples was determined by tapping mode. The sample scanning rate was between 1.2 and 2.2 Hz to minimize the imaging artifacts and tip-sample interaction.

## Acknowledgment

We thank Oguzhan Oguz for the help with the AFM imaging, and to Dr. Esra Yuca for her help with protein expression and purifications.

## Author Information

**Cemile Elif Özçelik**, UNAM-Institute of Materials Science and Nanotechnology, Bilkent University, Ankara, 06800, Turkey; Neuroscience Graduate Program, Bilkent University, Ankara, 06800, Turkey

**Özge Beğli**, UNAM-Institute of Materials Science and Nanotechnology, Bilkent University, Ankara, 06800, Turkey

**Ahmet Hınçer**, UNAM-Institute of Materials Science and Nanotechnology, Bilkent University, Ankara, 06800, Turkey

**Recep Erdem Ahan**, UNAM-Institute of Materials Science and Nanotechnology, Bilkent University, Ankara, 06800, Turkey

**Mehmet Seçkin Kesici**, Department of Chemistry, Faculty of Science, Middle East Technical University, Ankara 06800, Turkey

**Talip Serkan Kasırga**, UNAM-Institute of Materials Science and Nanotechnology, Bilkent University, Ankara, 06800, Turkey

**Salih Özçubukçu**, Department of Chemistry, Faculty of Science, Middle East Technical University, Ankara 06800, Turkey

**Urartu Özgür Şafak Şeker,** UNAM-Institute of Materials Science and Nanotechnology, Bilkent University, Ankara, 06800, Turkey; Neuroscience Graduate Program, Bilkent University, Ankara, 06800, Turkey

## Author Contributions

UOSS, conceived the idea; CEO, UOSS, OB designed the experiments, analyzed the data wrote the manuscript, OB prepared S. cerevisiae host cells for surface display, REA,AH, CEO carried out protein expression and purification studies; MSK and SO, synthesized the peptides; SK, carried out AFM experiments, SK and CEO analyzed the AFM data.

