## Supplementary file for "Synergistic Screening of Peptide-Based Biotechnological Drug Candidates for Neurodegenerative Diseases using Yeast Display and Phage Display"

**Supplementary Materials**

Cemile Elif Özçelik^1^, Özge Beğli^1^, Ahmet Hınçer^1^, Mehmet Seçkin Keskin^2^, Oğuzhan Oğuz^1^, Recep Erdem Ahan^1^, Talip Serkan Kasırga^1^, Salih Özçubukçu^2^, Urartu Özgür Şafak Şeker ^1,3^

^1^ UNAM- Institute of Materials Science and Nanotechnology, Bilkent University, Ankara, 06800, Turkey

^2^ Department of Chemistry, Faculty of Science, Middle East Technical University, Ankara 06800, Turkey

^3^ Neuroscience Graduate Program, Bilkent University, Ankara, 06800, Turkey

**Contents**

*Figure S1*

*Figure S2*

*Figure S3*

*Figure S4*

*Figure S5*

*Figure S6*

*Supplementary Methods*

*Supplementary References*

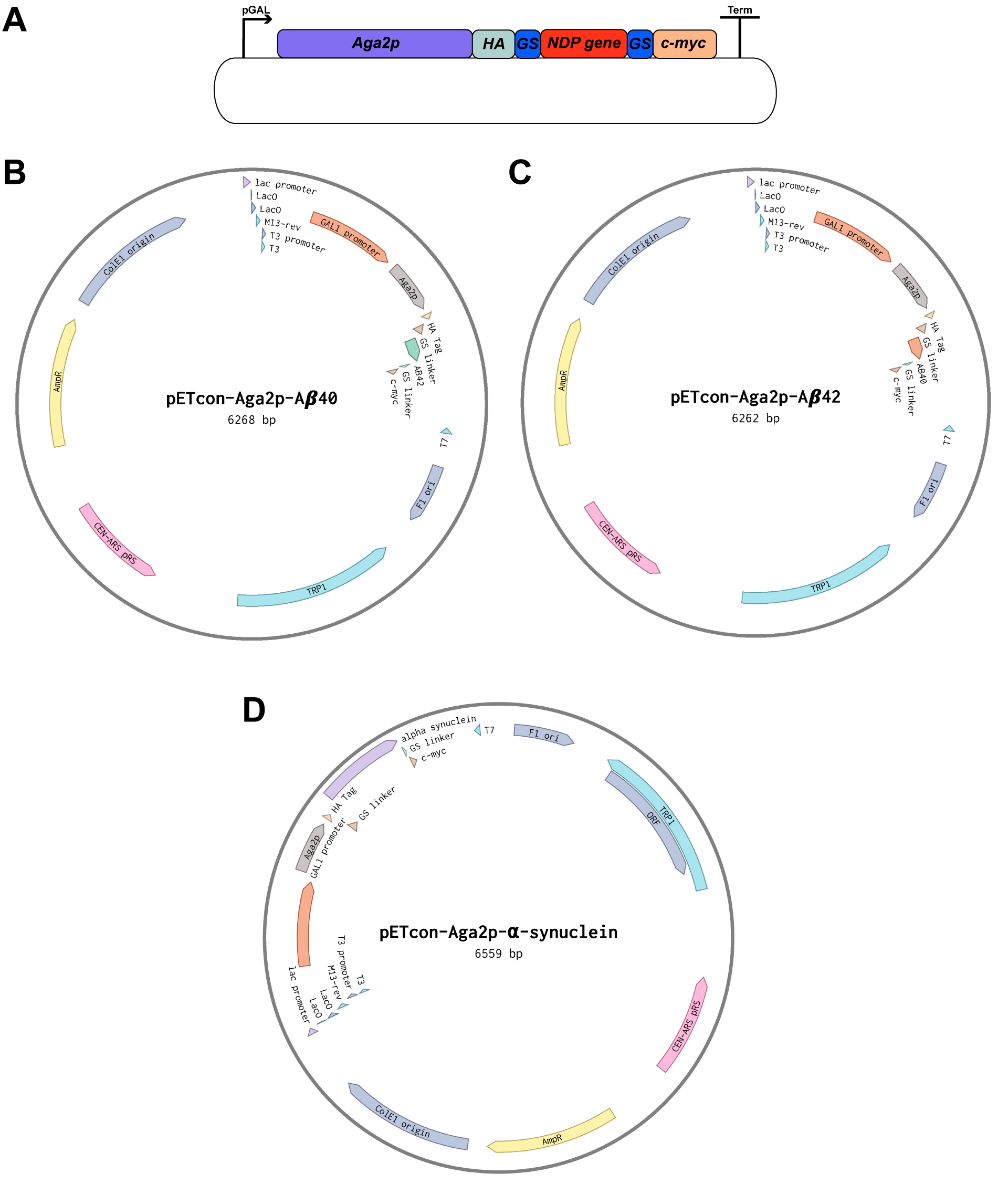

Figure S1. The representation of yeast surface display expression cassette. (A) All NDPs were expressed as a fusion of *Aga2p* gene under galactose-inducible promoter, pGAL. Aga2p and NDP gene were separated by GS linker. HA and c-myc existed as tags. c-myc tag were used for ICC experiments (B-D) The plasmid map for pETcon-Aga2p-Amyloid β_40_ was pETcon-Aga2p-Amyloid β_42_, and pETcon-Aga2p-⍺-synuclein plasmid maps were obtained from Benchling^1^.

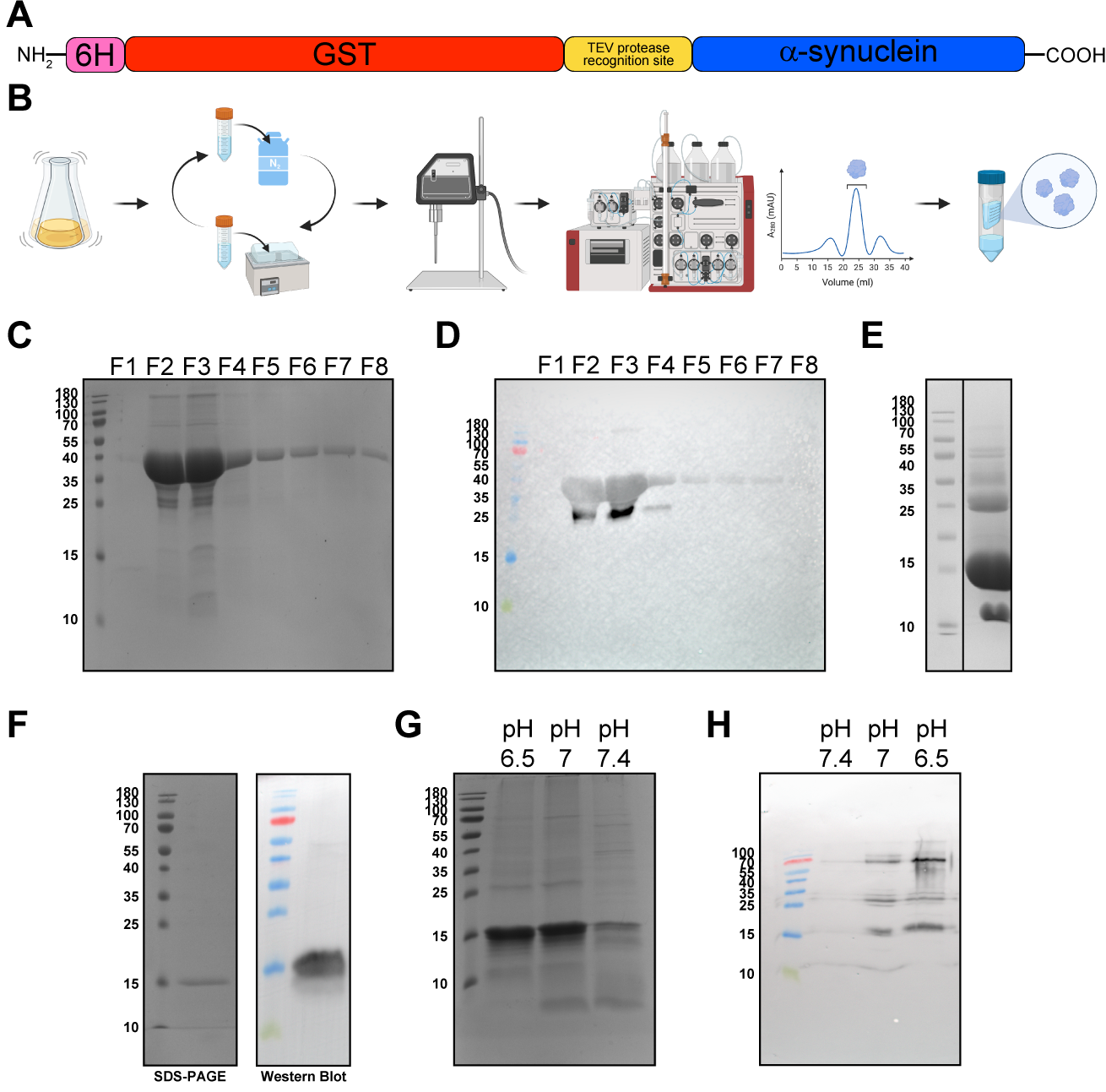

Figure S2. ⍺-synuclein monomers and fibrils were prepared by several FPLC method, and analyzed with SDS-PAGE and western blotting. (A) The design of ⍺-synuclein for protein purification. 6H was used for FPLC method, and GST tag was used for solubilization during expression and purification steps. (B) Workflow for protein purification was represented schematically. The overnight induced cells were lysed chemically that was followed by mechanical lysing with freeze-thaw cycles and sonication. After lysis, FPLC was used for protein purification. Purified proteins were concentrated for TEV protease reaction. For monomeric purification, FPLC step and protein concentration steps were repeated. The figure was created with [BioRender.com](https://biorender.com/) (C) The fractions obtained after 6H-GST-⍺-synuclein protein purifications were analyzed by SDS-PAGE. 40 kDa 6H-GST-⍺-synuclein were obtained in high concentrations in F2 and F3 fractions. (D) Western blotting was done with the same fractions obtained after 6H-GST-⍺-synuclein purification. (E) Western blotting was done after TEV reaction for cleave 14 kDa monomeric ⍺-synuclein from 6H-GST-⍺-synuclein. Removed GST and TEVp were observed around 26 kDa and 55kDa, respectively. (F) SDS-PAGE and western blotting were done for detection of monomeric ⍺-synuclein. (G-H) For the optimization of fibrillization pH, fibrillization assay samples with different pH values were analyzed SDS-PAGE. The monomeric ⍺-synuclein band intensities for samples from PB, pH 6.5 and PB, pH 7 were high, still there were low intensity bands in the upper parts of the gel. (H) Fibrillization assay samples with different pH values were analyzed by western blotting. Small sized seeds were detected well in the samples obtained with PB, 6.5 pH.

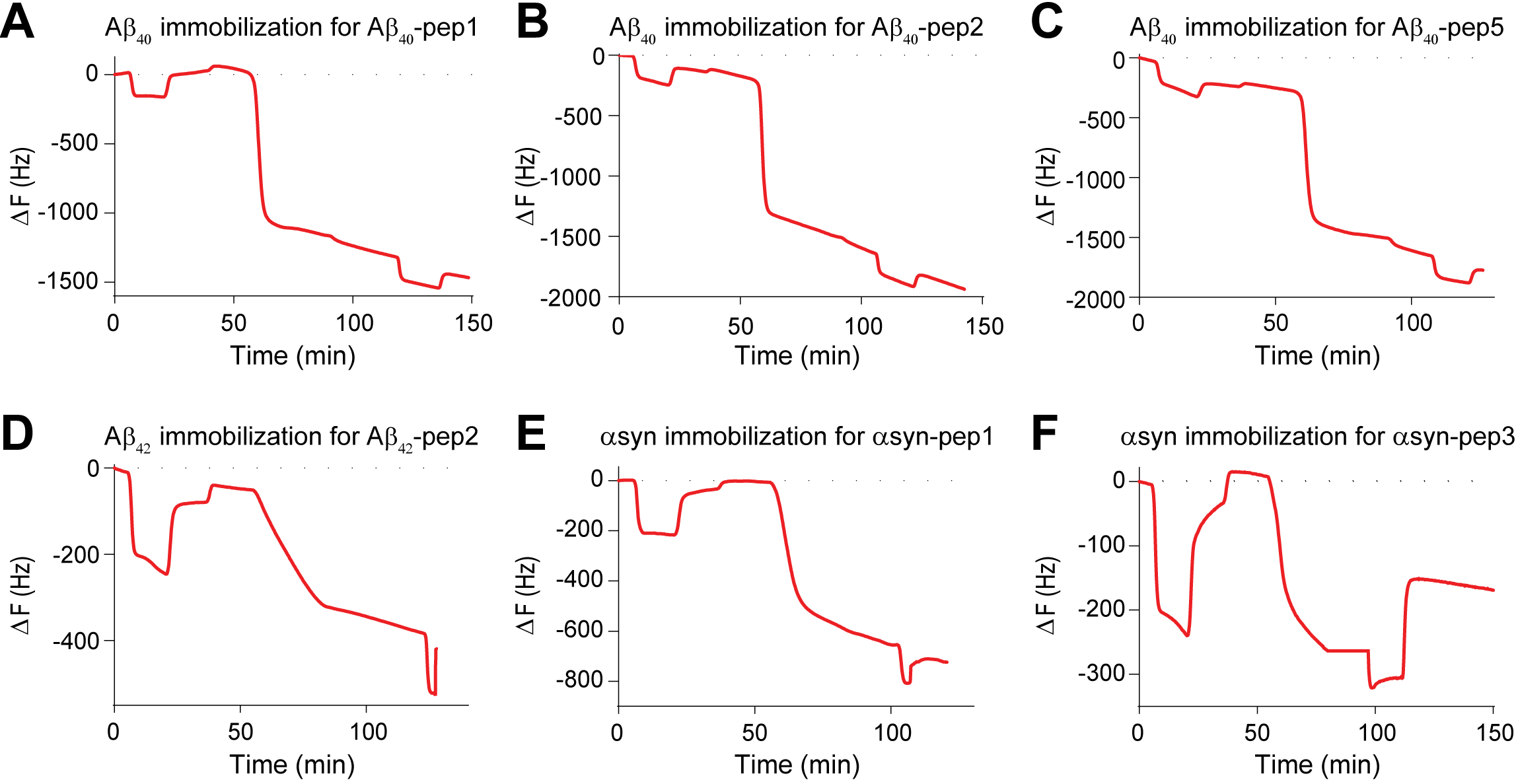

Figure S3. NDP-peptide analysis for QCM was achieved for each peptide by immobilizing each NDPs to gold chip by EDC/NHS coupling reaction. (A-C) Immobilization of amyloid β_40_ for Aβ_40_-pep1, Aβ_40_-pep2, and Aβ_40_-pep5 interactions were achieved with the mass depositions of 4536 ng.cm^-2^, 5090 ng.cm^-2^, 4648 ng.cm^-2^ on the chips, respectively. (D) Immobilization of amyloid β_42_ for Aβ_42_-pep2 interaction was achieved with the mass deposition of1106 ng.cm^-2^. (E-F) Immobilization of ⍺-synuclein for ⍺syn -pep1, and ⍺syn-pep3 interactions were achieved with the mass depositions of 1778 ng.cm^-2^, and 536 ng.cm^-2^ on the chips, respectively.
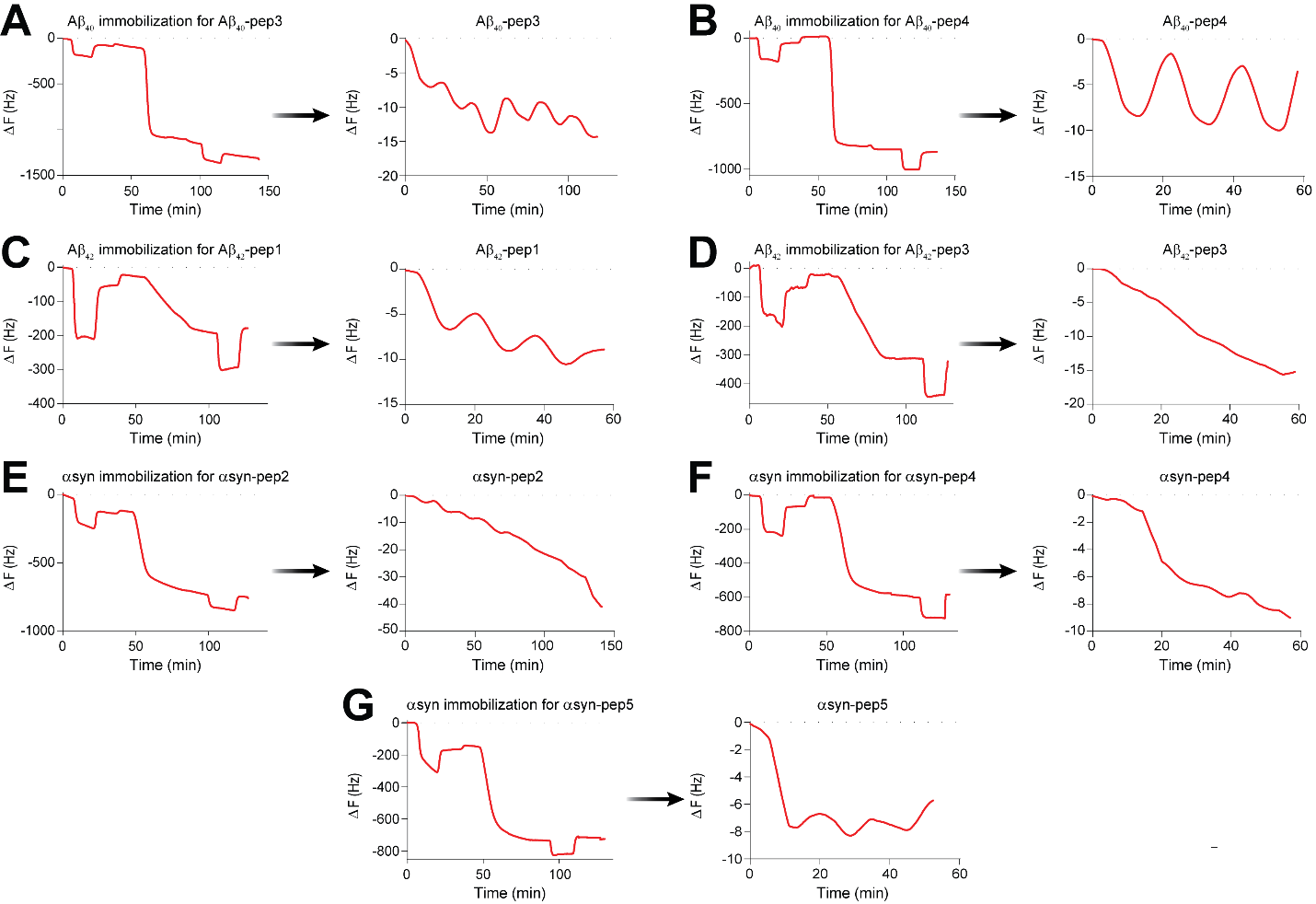

Figure S4. For all different peptides for NDPs were analyzed by QCM to determine protein-peptide interaction. First, gold chips were coated with the target NDP monomers by EDC/NHS coupling reaction. Then, peptides were introduced individually to detect whether interaction was occurred or not. (A) Immobilization of amyloid β_40_ for Aβ_40_-pep3 was achieved with the mass depositions of 3661 ng.cm^-2^ on the chips. The mass deposition of Aβ_40_-pep3 onto amyloid β_40_ coated gold chip was 405 ng.cm^-2^ (B) Immobilization of amyloid β_40_ for Aβ_40_-pep4 was achieved with the mass depositions of 2711 ng.cm^-2^ on the chips. The mass deposition of Aβ_40_-pep4 onto amyloid β_40_ coated gold chip was 9.5 ng.cm^-2^ (C) Immobilization of amyloid β_42_ for Aβ_42_-pep1 was achieved with the mass depositions of 2723 ng.cm^-2^ on the chips. The mass deposition of Aβ_42_-pep1 onto amyloid β_42_ coated gold chip was 134 ng.cm^-2^.(D) Immobilization of amyloid β_42_ for Aβ_42_-pep3 was achieved with the mass depositions of 856 ng.cm^-2^ on the chips. The mass deposition of Aβ_42_-pep3 onto amyloid β_42_ coated gold chip was 100 ng.cm^-2^.(E) Immobilization of ⍺-synuclein for ⍺syn -pep2 was achieved with the mass depositions of 1952 ng.cm^-2^ on the chips. The mass deposition of ⍺syn -pep2 onto ⍺-synuclein coated gold chip was 427 ng.cm^-2^ (F) Immobilization of ⍺-synuclein for ⍺syn -pep4 was achieved with the mass depositions of 1742 ng.cm^-2^ on the chips. The mass deposition of ⍺syn -pep4 onto ⍺-synuclein coated gold chip was 27.2 ng.cm^-2^ (G) Immobilization of ⍺-synuclein for ⍺syn -pep5 was achieved with the mass depositions of 1776 ng.cm^-2^ on the chips. The mass deposition of ⍺syn -pep5 onto ⍺-synuclein coated gold chip was 3.8 ng.cm^-2^.

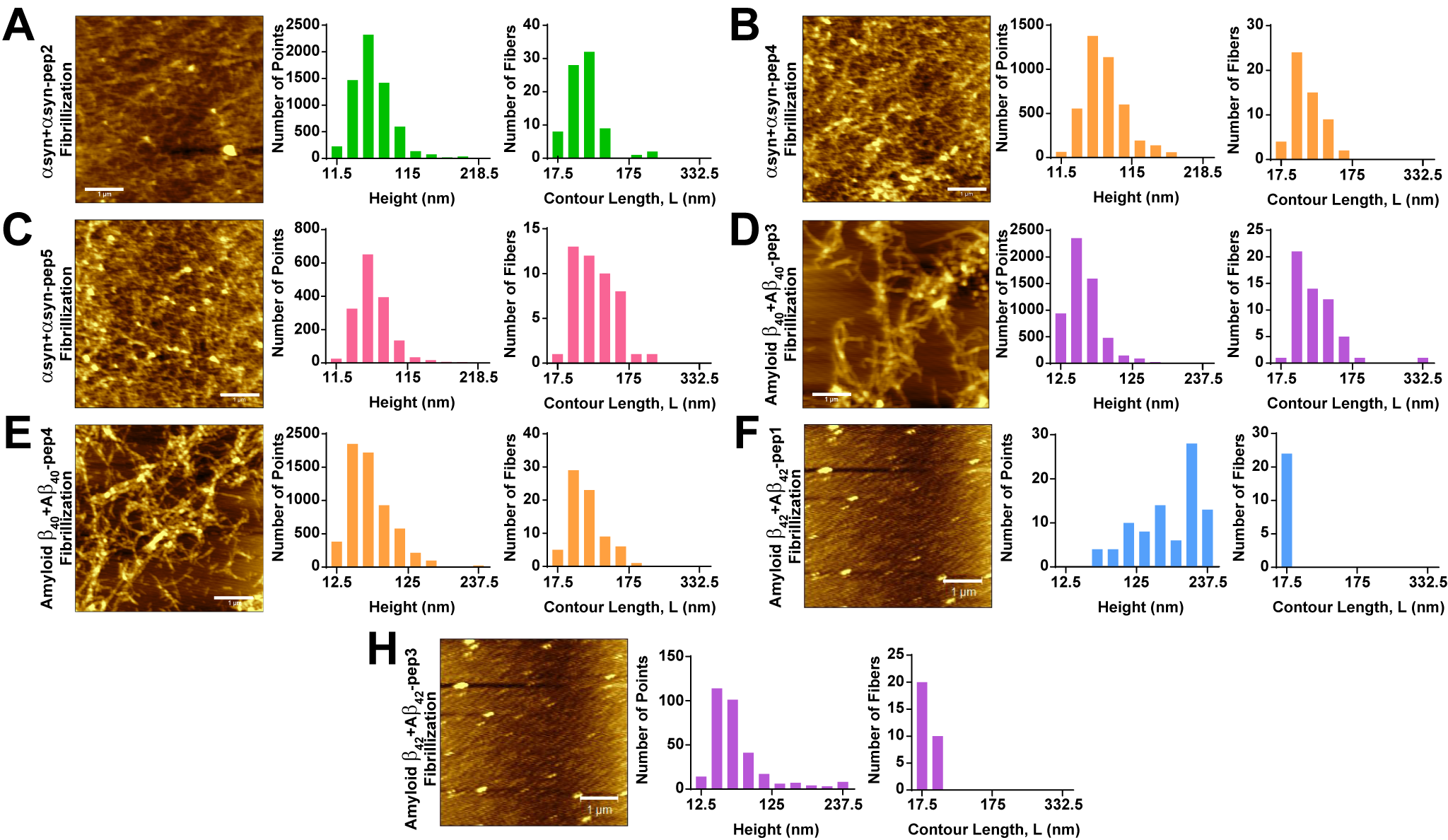

Figure S5. AFM analysis for fibrillization assay was applied to determine the effect of peptides in fibrillization. All fibril heights and lengths were analyzed by FiberApp (A) ⍺-synuclein+⍺syn -pep2 fibrillization product AFM result gave a result of web-like aggregation with fibrils. (B) ⍺-synuclein+⍺syn -pep4 fibrillization product AFM result gave a result of more intense web-like aggregation with fibrils. (C) ⍺-synuclein+⍺syn -pep5 fibrillization product AFM result gave a result of more web-like aggregation with fibrils with relatively large contour length. (D) Amyloid β_40_+Aβ_40_-pep3 fibrillization product AFM result gave a result of web-like aggregation with distinct fibrils. (E) Amyloid β_40_+Aβ_40_-pep4 fibrillization product AFM result gave a result of more intense web-like aggregation with shorter distinct fibrils.

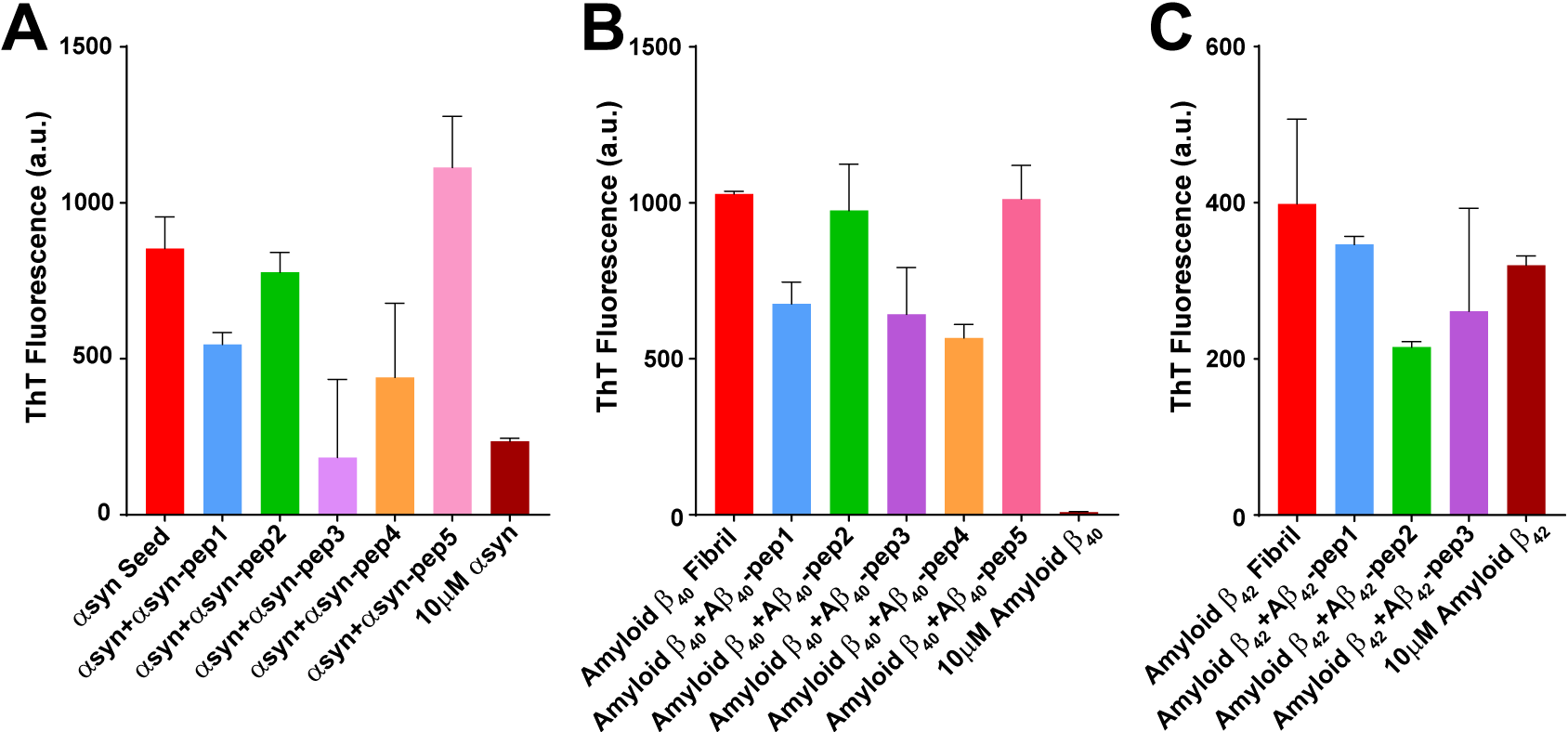

Figure S6. End point ThT fluorescence measurements gave different fluorescence signals for each protein-peptide interaction that were used to analyzed by AFM. (A) ThT fluorescence signals were measured for each fibrillization case of ⍺-synuclein. (B) ThT fluorescence signals were measured for each fibrillization case of amyloid β_40_. (C) ThT fluorescence signals were measured for each fibrillization case of amyloid β_42_.

**Supplementary Methods**

**Deprotection of resin and coupling with a Fmoc protected amino acid and cleavage reaction during solid phase peptide synthesis.** Rink Amide Resin (151.1 mg, 0.05 mmol) (Substitution: 0.331 meq/g) was weighed in a peptide synthesis reactor and it was washed with N, N-dimethylformamide (DMF), and the resin was swollen 20-30 min in 10 mL DMF. After the swelling part, 5 mL of 20% piperidine was poured into the reactor. After 3 min, this part was repeated a second time but 10 min. Then it was washed with DMF. After washing part, Fmoc protected amino acid (5.5 eq., 0.275 mmol) and HBTU (5.0 eq., 0.250 mmol) were weighed in a test tube, and 2.0 mL 0.3 M diisopropylethylamine (DIEA) in DMF was added. After the addition of the DIEA solution, the resulting mixture was added to the reaction vessel and the coupling mixture was left for 1 hour. After the completion of coupling, the resin was washed with DMF extensively. This process was repeated until the desired peptide elongation was obtained. At the end of peptide elongation and the last Fmoc deprotection, the resin was washed with DMF and DCM respectively and left for drying with the pump open for 30 min. After resin was dried, 5 mL cleavage cocktail, 95% TFA (4.75 mL), 5% Milli Q (0.125 mL) and 5% TIPS (0.125 mL) was prepared and added to the reactor. It was incubated for 2 hours and the cleaved peptide solution was precipitated with ice-cold diethyl ether, the solution was centrifuged and the peptide was washed three times with cold ether to remove small organic impurities.

**Thioflavin (ThT) Assay.** After fibrillization assay, 100 uM ⍺-synuclein and 10 uM peptide, 100 uM uM amyloid β_40_ and 250 uM peptide, 100 uM amyloid β_42_ and 250 uM peptide fibrillization products were added to 96-well plate with 1:10 dilution ratio in 1X PBS. Then NaN_3_ and 1M ThT in ddH_2_O were added onto each sample to final concentration as 0.1% and 25 uM, respectively. Samples were incubated at 37^o^C for 3 hours in dark. At the end of the incubation, fluorescence was measured at 450 nm excitation and 485 nm emission with 475 nm cutoff value by SpectraMax M5 microplate reader.
